# Impact of flushing procedures on drinking water biostability and invasion susceptibility in distribution systems

**DOI:** 10.1101/2025.02.19.639043

**Authors:** Fien Waegenaar, Thomas Pluym, Elise Vermeulen, Bart De Gusseme, Nico Boon

## Abstract

Maintaining microbial stability in drinking water distribution systems (DWDS) is a priority for water providers, yet operational events like flushing and the potential invasion of unwanted microorganisms pose significant challenges. This study utilized a pilot-scale DWDS to evaluate the susceptibility of drinking water networks after standard operational flushing procedures (with and without chlorination) as these procedures can alter biostability. Results showed that flushing with chlorination reduced bulk cell counts initially but led to subsequent microbial regrowth, highlighting the limitations of chlorination in long-term biostability. Biofilm cell densities and community composition were similar before and after flushing with and without chlorination. Additionally, the susceptibility of a flushed network against introduced bacterial indicators (*Aeromonas media*, *Pseudomonas putida*, and *Serratia fonticola*) was investigated. The invasion experiments revealed that the species decayed more rapidly in loops flushed with chlorination. This study emphasizes the trade-off between potential regrowth and microbial control, underscoring the need for strategies that enhance microbial resilience without fostering regrowth. These findings are critical for optimizing DWDS management practices to ensure drinking water safety and quality.

**Importance:** Flushing, with or without chlorination, is a standard procedure used by drinking water providers to address water quality issues such as brown water, as well as following waterworks, or contaminant detection. This process involves discharging large volumes of drinking water, often through fire hydrants. Depending on the contamination, additional disinfectants, such as free chlorine, are dosed during flushing. However, the longer-term effects of these processes on microbial dynamics are unknown. For example, while brown water issues can be solved, the overall biostability of drinking water may be compromised, potentially making the drinking water distribution systems more susceptible to the invasion of unwanted microorganisms. Studying these practices in full-scale distribution networks is challenging. Therefore, in this study, a unique drinking water distribution pilot system that mimics a real network with mature biofilm was used to investigate the changes in both bulk water and biofilm microbiomes following flushing procedures.

## 1 Introduction

Within the concept of drinking water biostability, the goal for water providers is to ensure a stable microbial community from the end of treatment to the consumer’s tap. Achieving biostability means establishing a microbial community that can resist environmental changes and unexpected events within drinking water distribution systems (DWDS)(1, 2). However, in practice, maintaining biostability poses significant challenges. Currently, biostability is often enforced by the addition of disinfection products(1, 3). Although, ozonation and chlorination for example, are leading to regrowth of naturally occurring microorganisms during distribution caused by the introduction of dead biomass and/or the depletion of the disinfection residual(4–8). This microbial regrowth can compromise water quality, leading to higher susceptibility to pathogens and/or undesirable changes in color, taste, or odor(1). Following quality issues such as brown water, as well as after waterworks, maintenance, or contaminant detection, the local distribution network is typically flushed by discharging large volumes of drinking water, often at fire hydrants. Depending on the contamination, additional disinfectants, such as free chlorine, are dosed during flushing, and customers are advised not to drink from the tap(9–13). These flushing procedures can mobilize materials accumulated on the pipe walls into the bulk water, leading to aesthetically unacceptable drinking water and/or release of unwanted microorganisms into the network(14–16). In a study by Douterelo et al.(10), a section of a full-scale network was flushed using hydrants at a velocity of 0.60 m/s. Analysis of the flushed water revealed elevated turbidity levels(10). Other studies used pilot-scale distribution systems, assessing flushing efficacy on young biofilms of 28 days (velocity: 0.57 m/s, shear: 0.89 N/m²)(17) or on pipe segments from the full-scale network with 12-month-old biofilms (velocity: 0.67 - 1.9 m/s, shear: 1.99 - 7.22 N/m²)(16). Both studies concluded that mechanical biofilm removal through flushing was insufficient to completely remove bacteria from the pipe walls. In addition, it is shown that shock-chlorination with the addition of 3.7 mg Cl_2_/L reduces biofilm density but also affects extracellular protein substances (EPS) contact points(18). Furthermore, additional chlorination can increase assimilable organic carbon (AOC) concentrations, thereby increasing the potential for microbial (re)growth in the DWDS(6). Consequently, flushing and shock chlorination are expected to have long-term impacts on the biostability of drinking water(19). However, studies investigating the longer-term effects of these processes on microbial dynamics remain scarce in the literature. For example, while brown water issues can be solved, the overall biostability of drinking water may be compromised, potentially making the DWDS more susceptible to the invasion of unwanted microorganisms.

Drinking water contamination during distribution can occur due to the infiltration of contaminants during water works, or accidental connections with non-potable water sources. On the other hand, biofilms can harbor pathogens or other unwanted microorganisms, which can be released into the water after for example flushing procedures(20, 21). Additionally, opportunistic pathogens present in the DWDS, even if they are below detection limits, can proliferate due to nutrient introduction from dead cells or biofilm shear stress(4). Previous studies have detected pathogens, such as *Pseudomonas aeruginosa*, *Mycobacteria* spp., and *Legionella* spp., as well as fecal indicators, such as *Escherichia coli*, in biofilm samples from full-scale distribution networks(22–24). To control and evaluate drinking water contamination, the measurement of *Escherichia coli* and coliform bacteria is the standard parameter in Europe, with the requirement being absent in 100 mL samples. The coliform *Serratia fonticola* is often detected in the Flemish distribution network (Belgium)(13, 25–27). Although it struggles to survive in the oligotrophic drinking water environment, the coliform is able to attach and get established within the mature biofilms(26, 28, 29). Furthermore, not only indicator bacteria for fecal contaminations are regulated, but also pathogens such as *Pseudomonas aeruginosa* and *Clostridium perfringens*, which must be absent in 250 mL and 100 mL samples, respectively(13, 30). Previous studies have shown that *Pseudomonas aeruginosa* is often associated with the drinking water biofilm and exhibits high resistance to chlorine(24, 31–34). Other non-pathogenic *Pseudomonas* spp., though not regulated, are often linked to external contaminations like rainwater intrusions due to improper connections(24, 30, 35). To control microbial regrowth, *Aeromonas* spp. are measured and included in the Dutch Drinking Water Decree as an indicator for elevated microbial regrowth in non-chlorinated DWDS(36). The legal standard for *Aeromonas* spp. in drinking water is 1000 CFU/100 mL. In the Netherlands, *Aeromonas rivuli*, *Aeromonas veronii*, *Aeromonas sobria*, and *Aeromonas media* are mostly detected in the DWDS(37). Previous studies have mainly focused on the detection of unwanted microorganisms rather than on the growth and survival dynamics within the oligotrophic drinking water ecosystem and in pretreated networks(37, 38). Though, understanding growth dynamics is important, as the potential for invasion is depending on nutrient availability and competition with the indigenous drinking water community(39, 40).

Here, we used a pilot-scale DWDS with a mature biofilm to study the bacterial bulk water and biofilm communities. The goal was to evaluate the susceptibility of drinking water networks after standard operational flushing procedures as these procedures can alter biostability. First, we investigated the direct and long-term impact of flushing with and without chlorination on drinking water the bulk and biofilm microbiome. Secondly, the survival of three unwanted microorganisms (i.e., *Aeromonas media*, *Pseudomonas putida*, *Serratia fonticola*) was evaluated, and the susceptibility of the biofilms to these bacterial indicators was evaluated. Membrane filtration in combination with selective plating was used to monitor the dynamics of the unwanted microorganisms. Bulk water quality was monitored using online flow cytometry and 16S rRNA gene-base amplicon sequencing. Additionally, a coupon system was used to sample and analyze biofilm cell densities and its corresponding community composition.

## 2 Materials and Methods

### 2.1 Experimental design

A DWDS pilot was used as described in García-Timermans et al.(41) (Fig. A1). Briefly, the pilot consists of three identical subsystems (i.e., loops). Each loop has a 1 m³ HDPE intermediate bulk container (IBC) connected to 100 m of 80 mm PVC-U pipes. Water is pumped from the IBC, recirculates through the loops, and returns. The pilot was fed with tap water from Farys (Ghent, Belgium), sourced from the Albert Channel in Antwerp.

### 2.2 Flushing experiment

First, a flushing experiment was conducted to investigate the influence of standard operational procedures performed by water providers in cases of contamination. Depending on the contamination, additional disinfectants, such as free chlorine, are dosed during flushing. Loop 1 was flushed with tap water and NaOCl (0.5 mg/L free chlorine), followed by two flushes with tap water only. Loop 2 (control) was not flushed, while loop 3 underwent three flushes with tap water (Table 1). In detail, each loop received 650 L of tap water. For loop 1, after one cycle at 150 L/min, 33.07 mL of NaOCl (14% Cl₂, Avantor) was added to the IBC to reach 0.5 mg/L free chlorine. Free chlorine was measured before, during, and after dosing to ensure uniform distribution. Upon stabilization of the concentration, flushing was conducted at 150 L/min for 4.3 minutes (i.e., 0.5 m/s). The system was then drained and refilled, repeating the flush twice without NaOCl. Loop 3 followed the same protocol without NaOCl, while Loop 2 recirculated water for ∼7 hours at 24 L/min. Bulk samples for flow cytometry, 16S rRNA gene-based amplicon sequencing, ATP, TOC, and free chlorine measurements were collected before, during, and after the flushing processes (section 2.5, 2.7) following WAC/I/A/001(42). Biofilm samples (n = 3) were taken before and after flushing (section 2.6). This flushing experiment was performed twice, with a two-month interval between each.

**Table 1:**
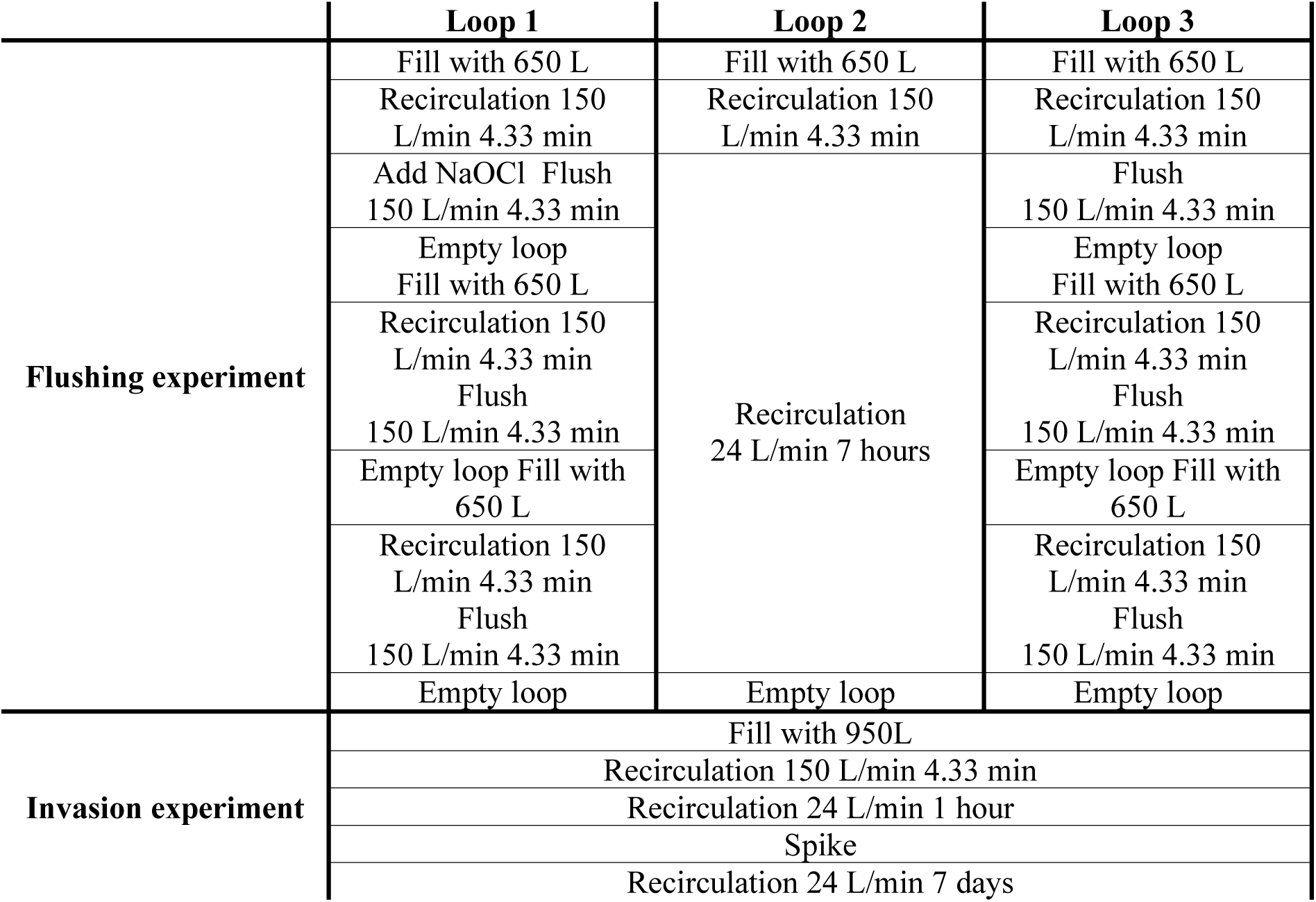
Experimental set-up of the flushing and invasion experiment for each loop. The experiments were conducted twice, once at 16°C and once at 20°C.

### 2.3 Invasion experiment

After each flushing experiment, loops were refilled with 950 L of tap water and recirculated for 1 hour at 24 L/min before sampling for 16S rRNA sequencing, ATP, TOC, and free chlorine. (Table 1). Subsequently, 1 mL of a 10⁶ cells/mL solution of unwanted microorganisms was added to each loop, and survival was monitored for 7 days via selective plating (section 2.4). The experiment ran twice (16°C and 20°C) with a 7-day hydraulic residence time. In the first experiment, 500 L was replaced twice weekly, while in the second experiment, all water was replaced after 7 days. Throughout the experiment, the flow velocity was 24 L/min, and the pressure was kept between 0.7 and 0.9 bar(14). Conductivity, flow velocity, pressure, pH, and temperature were logged every 5 minutes(41). Online flow cytometry measured total cell densities every 8 (experiment 16°C) or 12 (experiment 20°C) hours. Bulk water and biofilm coupons (n = 3) were analyzed for microbial and chemical parameters. Bulk water samples were collected for microbial (ATP, 16S rRNA gene-based amplicon sequencing) and TOC analysis (section 2.5, 2.7). Biofilm coupons (n = 3) were taken after 7 days and further analyzed (section 2.6).

### 2.4 Bacterial indicators and culture conditions

The bacterial indicators, *Aeromonas media*, *Pseudomonas putida*, and *Serratia fonticola*, were isolated from the Flemish drinking water distribution network (Pidpa in Antwerp, De Watergroep in East-West Flanders, and Farys in Ghent, respectively) and identified with 16S rRNA gene Sanger sequencing as described by Kerckhof et al.(43). Strains were revived from the −80 °C stock, streaked on a Reasoner’s 2A (R2A) (18.1 g/L final concentration) agar plates (Oxoid, England) and incubated at 28°C for 24 hours. Colonies were resuspended in 5 mL R2A broth (3 g/L, Oxoid, England) and incubated (28°C, 100 rpm, 24 h). Cultures were washed three times with sterile 8.5% NaCl, centrifuged (2500 × g, 5 min), then transferred to diluted R2A broth (50 mg/L) for another 24 h incubation. After repeating the washing steps, cells were resuspended in 0.2 µm filtered sterile 8.5% NaCl for intact cell count (ICC) measurement via flow cytometry. Cultures were diluted to 10⁶ cells/mL, and 1 mL was added per loop, achieving spike concentrations of 100-600 cells/100 mL.

### 2.5 Microbial monitoring of the bulk water phase

Online flow cytometry measured total cell densities and performed phenotypic fingerprinting using an onCyt© autosampler (onCyt Microbiology AG, Switzerland) coupled to an Accuri™ C6 Plus flow cytometer (BD Biosciences, Belgium) as described in Waegenaar et al.(44) (Fig. A1). Briefly, samples (200 µL) were taken in triplicate for each loop every 4 hours (experiment 16°C) or every 8 hours (experiment 20°C), stained with SYBR Green I (5000× diluted in TRIS buffer, pH 8), incubated at 37°C for 20 min, and analyzed. The onCyt sample lines were cleaned with sodium hypochlorite (1 v% final concentration, Avantor, USA), neutralized with sodium thiosulfate (50 mM final concentration, Merck, Belgium), and rinsed with ultrapure water (Milli-Q, Merck, Belgium). Control samples and samples collected during the flush experiment were manually collected and measured with similar staining and incubation conditions on an Accuri^TM^ C6 Plus flow cytometer (BD Biosciences, Belgium) in the lab.

Before, during, and after the flushing processes, and after 7 days, samples for 16S rRNA gene-based amplicon sequencing were taken. From each loop, 2 L was filtered (0.22 μm MCE Membrane filter (Merck, Belgium)) using a filtration unit consisting of six filtration funnels and a Microsart e.jet vacuum pump (Sartorius, Germany), stored in a freezing tube at −21°C, and processed as per Waegenaar et al.(44). Briefly, DNA extraction was performed using the DNeasy PowerSoilPro kit (Qiagen, Germany), and 10 μL genomic DNA extract was send out to LGC genomics GmbH (Berlin, Germany) for library preparation and sequencing on an Illumina Miseq platform with v3 chemistry (Illumina, USA).

Bacterial indicator concentrations in bulk water were determined by filtering dilutions (3 × 100 mL) through 0.45 µm S-Pack filters (Merck, Belgium) using a filtration unit and a Microsart e.jet vacuum pump (Sartorius, Germany), incubating at 37°C for 18-24 h on selective agar: *Aeromonas media* (Ampicillin Dextrin Agar (ADA), HiMedia, Germany), *Pseudomonas putida* (Pseudomonas Cetrimide (PCN) Agar, VWR, Netherlands), and *Serratia fonticola* (Chromogenic Coliform Agar (CCA), Carl Roth, Belgium), following ISO 9308-1:2014(45). For PCN agar, 15 mL glycerol (≥99.5%, Carl Roth, Belgium) was added per 1000 mL before autoclaving (121°C). For ADA, ampicillin (HiMedia, Germany) was added after cooling to ∼50°C. Bulk water was filtered before spiking for controls. Samples were collected at multiple timepoints after the spike (0.33 to 304 h). Random selective plates were confirmed using matrix-assisted laser desorption-ionization–time of flight mass spectrometry (MALDI-TOF MS) (Vitek MS, bioMérieux, Marcy-l’Étoile, France).

### 2.6 Biofilm sampling

Biofilm was sampled using PVC-U coupons installed in each loop (Fig. A1C, D) and cultivated for one year. Coupons (three for 16°C, two or three for 20°C) were taken before and after flushing and after 7 days. Biofilm cells were removed using an electric toothbrush (Oral-B, Advanced Power) into 15 mL of 0.2 μm filtered bottled water (Evian, France) as described in Waegenaar et al.(44). The biofilm suspensions (10× diluted in 0.2 µm filtered bottled water (Evian, France)) were measured with flow cytometry using an Attune NxT BVXX flow cytometer (ThermoFisher Scientific, USA). For both experiments, staining was done with 1 v% of 100 times diluted SYBR Green I (10000× concentrate in 0.22 µm-filtered DMSO, Invitrogen, Belgium) solution to measure total cell counts(27). For the second experiment (20 °C), staining was also done with 1 v% of 100 times diluted SYBR Green I combined with propidium iodide (10000× concentrate in 0.22-µm filtered DMSO, 50 × 20 mM propidium iodide in 0.22-µm filtered DMSO, Invitrogen, Belgium) to measure intact-damaged cells. Incubation was at 37°C for 20 min in the dark, and measurements were in quadruplicate. Additionally, 3 mL (3×) of each suspension was filtered (0.45 µm S-Pack filters, Merck) for selective plating of *Aeromonas media*, *Pseudomonas putida*, and *Serratia fonticola* (section 2.4). The remaining volume was filtered (Millipore Express PLUS Membranes (Merck, Belgium), Polycarbonate syringe filter holder (Sartorius, Germany)) for 16S rRNA gene-based amplicon sequencing (section 2.5).

### 2.7 Free chlorine and total organic carbon (TOC) measurements

Free chlorine concentrations were determined using a Pocket Colorimeter II (Hach, Belgium) (detection limit = 0.02 mg/L). Samples for TOC analyses were collected in 40 mL TOC-free vials (Sievers, Germany) and stored at 6°C prior to analysis. TOC concentrations were measured in technical triplicate using a Sievers 900 Portable TOC Analyzer connected to a Sievers 900 Inorganic Carbon Remover (General Electric Company, Boston, USA).

### 2.8 Data analysis

Data analysis was done in R(46) RStudio version 4.3.0(47). The Flow Cytometry Standard files were imported using the flowCore package (v2.14.0)(48). The background data was removed by manually drawing a gate on the FL1-H (green) and FL3-H (red) fluorescence channels as described in Props et al.(49). Illumina data was processed using the DADA2 pipeline (v1.30.0)(50). Taxonomy was assigned using the Silva database v138(51). Normalization of the sample reads was done to correct for differences in sequencing depth among samples. The sequencing reads were ranging from 1017 to 101880. Further data analysis was performed using packages such as the phyloseq package (v1.46.0) and the vegan package (v2.6-4)(52, 53). The data generated by MALDI-TOF MS was analyzed using the MYLA® software (Pidpa, Antwerp). Data visualization was done using the ggplot2 (v3.4.4) and ggpubr (v0.6.0) packages(54, 55). Shapiro-Wilk Test was used to test the data for normality and further statistical analysis was done with the dplyr package (v1.1.4) and the vegan package (v2.6-4)(53, 56). In all cases, numbers following the ± sign are standard deviations (s.d.). To evaluate the decay of each unwanted microorganism during the invasion experiments, first-order decay rate constants (k (h^−1^)) were calculated as the slope of the line when ln(C_t_/C_0_) was regressed against time (t), where C_t_ is the concentration of the concerned microorganism (CFU/100 mL) at a certain time t and C_0_ is the concentration of the concerned microorganism (CFU/100 mL) at time 0(57). In the first experiment, loop 2 was not completely mixed after 20 minutes, so 1 hour was chosen as time 0 for this loop. In the second experiment, loop 3 was not completely mixed after 20 minutes, so 1 hour was selected as time 0 for this loop.

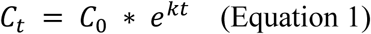

Shear stress values (*τ*) were calculated using Equation 2 for turbulent conditions (Reynolds number > 4000). The density (*ρ*) of water is 1000 kg/m³, the velocity (*u*) is calculated using the cross sectional area of a pipe and the flow velocity during the flushing process.

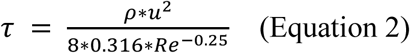

## 3 Results

### 3.1 Impact of flushing with or without chlorination on the bulk water phase

The impact of flushing with and without chlorination on the drinking water bulk and biofilm microbiome was studied using a pilot-scale distribution system with 3 identical loops. Loop 1 was flushed with water containing sodium hypochlorite (NaOCl), resulting in a free chlorine concentration of ± 0.50 mg HOCl/L, followed by two flushes with tap water without the addition of NaOCl, loop 2 served as a control and was not flushed, while loop 3 underwent three flushing cycles (Table 1). This experiment was performed twice. To evaluate the susceptibility of a system after flushing, an invasion experiment was performed after each flushing experiment, once at 16°C and once at 20°C. The free chlorine concentrations in the tap water supplied to each loop, as well as after the first flush of each loop, were below the detection limit of 0.02 mg Cl_2_/L. During each flushing experiment, samples were taken to measure TOC concentrations and total cell counts. The flush with chlorination led to a 1-log reduction in cell counts and an increase in TOC concentrations from 2.54 ± 0.15 mg/L to 4.43 ± 0.02 mg/L during the first flush of loop 1 (Fig. A2). After this flush, both the cell densities and TOC concentrations returned to levels similar to the other loops (± 3 × 10⁴ cells/mL and 2 mg TOC/L). Following the flushing experiment and during the invasion experiment at 20°C, an increase in TOC content up to 5.10 ± 0.14 mg/L was observed in loop 3 (Fig. A2).

Before, after the flush and during the invasion experiment, online flow cytometry was used to measure bacterial total cell densities and phenotypic traits of the community. Before the first flush experiment, the cell density in all loops followed the same increasing trend because of recycling the water for 7 days (Fig. 1A). On the day of the experiment (day 0), the initial cell density in loop 1, 2 and 3, was (2.76 ± 0.07) × 10^4^ cells/mL, (3.53 ± 0.16) × 10^4^ cells/mL and (3.53 ± 0.12) × 10^4^ cells/mL respectively. After 2 days, loop 1 exhibited a higher bacterial cell concentration compared to loop 2 and 3 and this increase was further observed after each refreshment. The maximum cell density was (3.75 ± 0.03) × 10^5^ cells/mL for loop 1, compared to (2.02 ± 0.08) × 10^5^ cells/mL and (1.58 ± 0.01) × 10^5^ cells/mL for loop 2 and 3, respectively. Statistical analysis showed a significant difference between the cell densities in loop 1 and the other loops (Dunn test with Holm correction, p_(loop 1 & 2)_ ≤ 0.05, p_(loop 1 & 3)_ ≤ 0.05), and no significant difference between loop 2 and 3 (p_(loop 2 & 3)_ = 0.19). This was further supported by ATP measurements, where after 7 days, loop 1 exhibited an ATP concentration of 50.27 ± 2.14 ng/L, compared to 6.75 ± 0.35 ng/L and 7.15 ± 0.40 ng/L for loop 2 and 3, respectively (Fig. A3). For the second flushing experiment, the cell density of loop 1 was already higher than the cell density of loop 2 and 3 before performing the flush (Fig. 1B). After the flush with chlorination, the bacterial cell concentration in loop 1 increased even more, up to (5.22 ± 0.17) × 10^5^ cells/mL, while loop 2 and 3 had similar lower cell densities and again a significant difference was observed between loop 1 and the other loops (Dunn test with Holm correction, p_(loop 1 & 2)_ ≤ 0.05, p_(loop 1 & 3)_ ≤ 0.05, p_(loop 2 & 3)_ = 0.37). However, the ATP concentrations were similar (i.e., ± 6.36 ng/L) between the loops before and after the flushing experiment (Fig. A3).

**Fig. 1:**
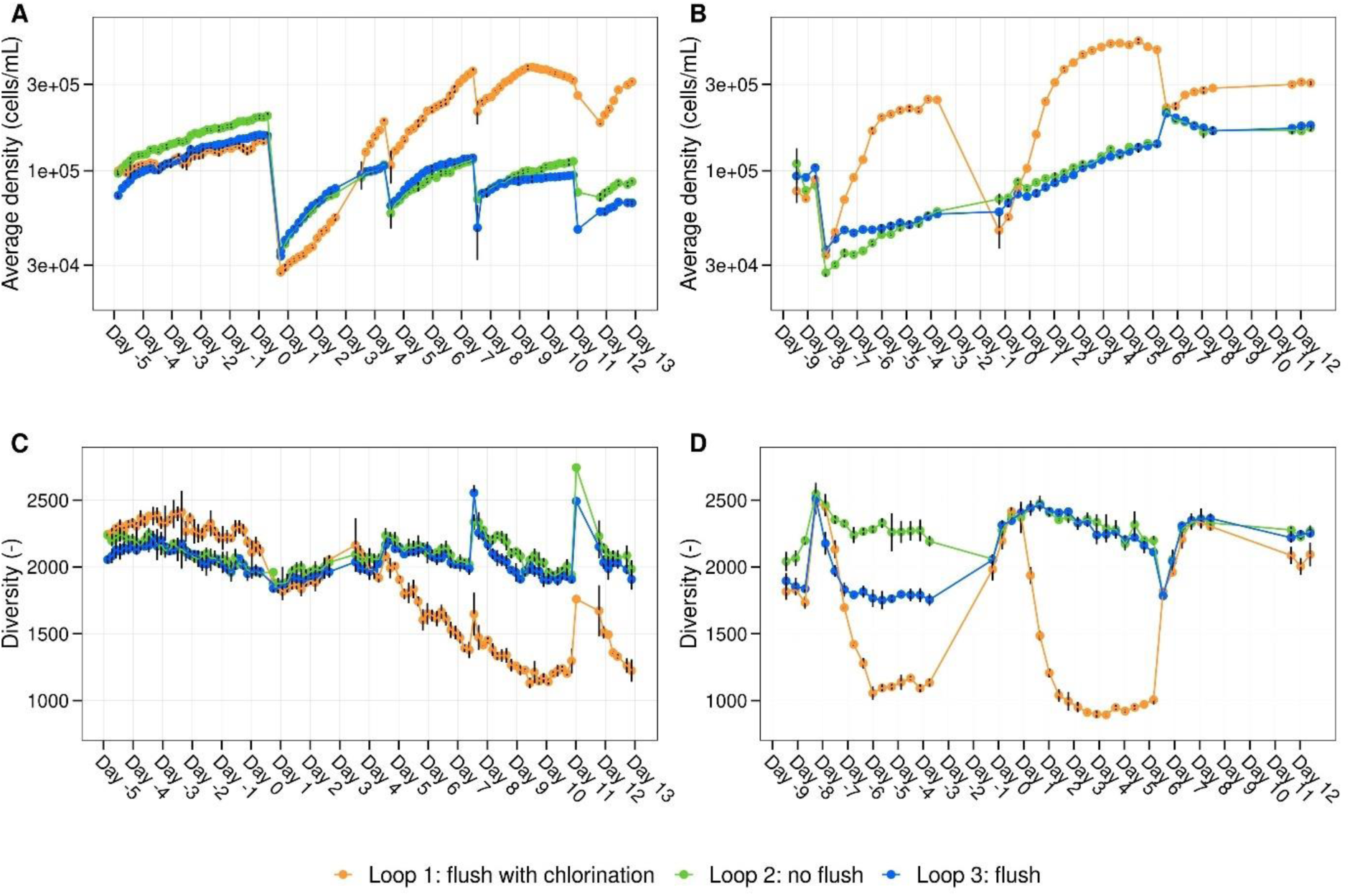
Cell density and phenotypic diversity of the bulk water community in function of time. Average density (cells/mL) in function of time (days) of the bacterial bulk community in loop 1 (orange), loop 2 (green), and loop 3 (blue) at **(A)** 16°C and **(B)** 20°C. Phenotypic diversity (D_2_) in function of time for each loop is shown in **(C)** and **(D)** for 16°C and 20°C, respectively. Day 0 represents the start of the experiment, where loop 1 underwent a flush with chlorine, loop 2 received no flushing, and loop 3 was flushed without chlorine. This flushing experiment was followed by an invasion experiment. Per timepoint, biological replicates (n = 3) were taken and corresponding error bars are shown in black.

In addition to the higher cell densities measured in loop 1, a lower phenotypic diversity was observed compared to loop 2 and 3 (Fig. 1C, D). This was further supported by 16S rRNA gene-based amplicon sequencing results, which revealed substantial growth of the family *Sphingomonadaceae*, specifically the genus *Sphingopyxis*, after 7 days in loop 1 (Fig. 2). *Sphingopyxis* accounted for 44.79% and 68.50% of the bacterial population in the experiments conducted at 16°C and 20°C, respectively. The presence of this genus may be attributed either to the tap water supplied to the pilot system (with relative abundances of 0.25% and 6.00% in experiments 1 and 2, respectively) or to interactions with the biofilm (with relative abundances in loop 1 of 0.77% and 7.15% in experiments 1 and 2, respectively).

**Fig. 2:**
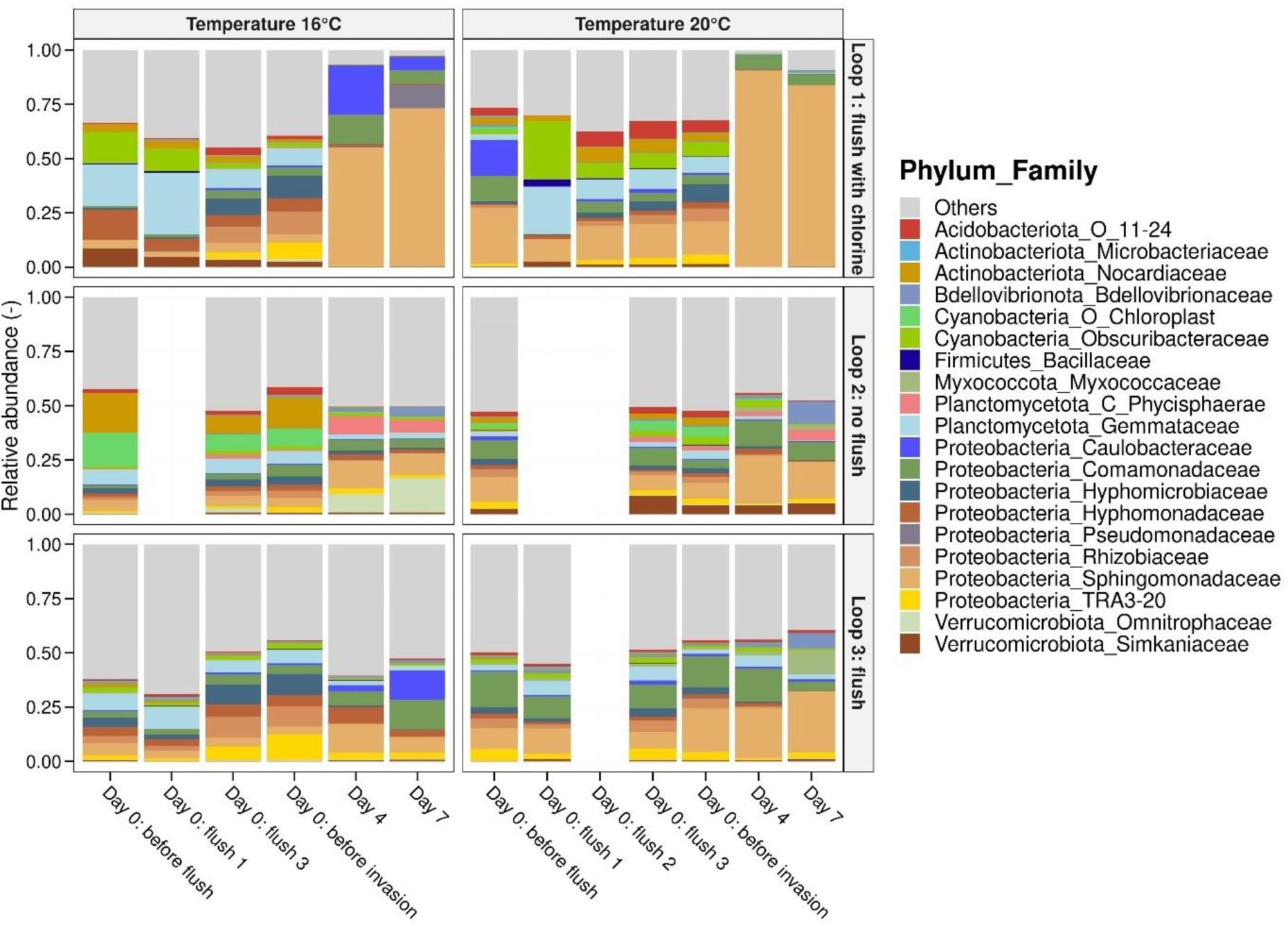
Bacterial community composition in the bulk water samples. Relative abundances of the 20 most abundant families of the bulk water before the flush, during the flush, after the flush and after 4 and 7 days. At each timepoint, one water sample was taken per loop (n = 1).

The bacterial composition of the flushed water samples appeared similar to that of the water before the flush, consisting mainly of *Proteobacteria* and *Actinobacteriota* (Fig. 2). In addition, Bray-Curtis dissimilarity values were calculated (Fig. A4). For flushed water samples without chlorine treatment, the Bray-Curtis dissimilarity values ranged between 0.27 and 0.41, indicating moderate similarity between these samples and the incoming water. On the other hand, the dissimilarity between the flushed water samples and the pre-flush water for loop 1 was higher, with values ranging from 0.32 to 0.85, showing more pronounced differences in bacterial composition and suggesting some degree of biofilm detachment. For loop 2, where the water recirculated for 7 hours at 24 L/min, the average dissimilarity index remained around 0.33, indicating minimal changes in bacterial communities. In this loop, bacteria from the *Chloroplast* order were detected, likely due to light exposure on the connection tube between the IBC and the pump.

### 3.2 Minimal impact of flushing with or without chlorination on the biofilm

To evaluate the effect of flushing on the drinking water biofilm, coupons composed of the same material as the pipes (PVC-U) were taken before and after the performed flushes (Fig. A1). The biofilm in the pilot system had been cultivated for one year, and the cell densities before the first flush experiment in loops 1, 2, and 3 were (5.40 ± 2.77) × 10^6^ cells/cm², (5.69 ± 3.90) × 10^6^ cells/cm² and (2.60 ± 0.74) × 10^6^ cells/cm² respectively, indicating the presence of a mature biofilm (Fig. A5A). The biofilm microbiome mainly consisted of *Proteobacteria* (± 50%), more specifically, *Xanthobacteraceae*, *Sphingomonadaceae*, *Rhodocyclaceae*, *Rhodobacteraceae*, and, *Comamonadaceae* and bacteria from the phylum *Dadabacteriales* (± 10 %), the phylum *Planctomycetota* (± 10 %), and *Verrucomicrobiota* (± 10 %) (Fig. 3). In both temperature scenarios, total cell densities remained consistent all loops before and after flushing (Fig. A5A). Moreover, no significant effect of flushing with or without chlorine was observed on the biofilm cell densities (Mann-Whitney test, p > 0.05) or bacterial composition (PERMANOVA, p > 0.05) (Fig. A5A, 3). In the second experiment, data on living and damaged cells was also collected, and an increase from (1.08 ± 0.14) × 10^6^ cells/cm² to (2.57 ± 0.26) × 10^6^ cells/cm² in damaged cells was observed in the biofilm of loop 1 after flushing and chlorination, whereas the number of damaged cells in the other loops remained constant before and after flushing (Fig. A5B).

**Fig. 3:**
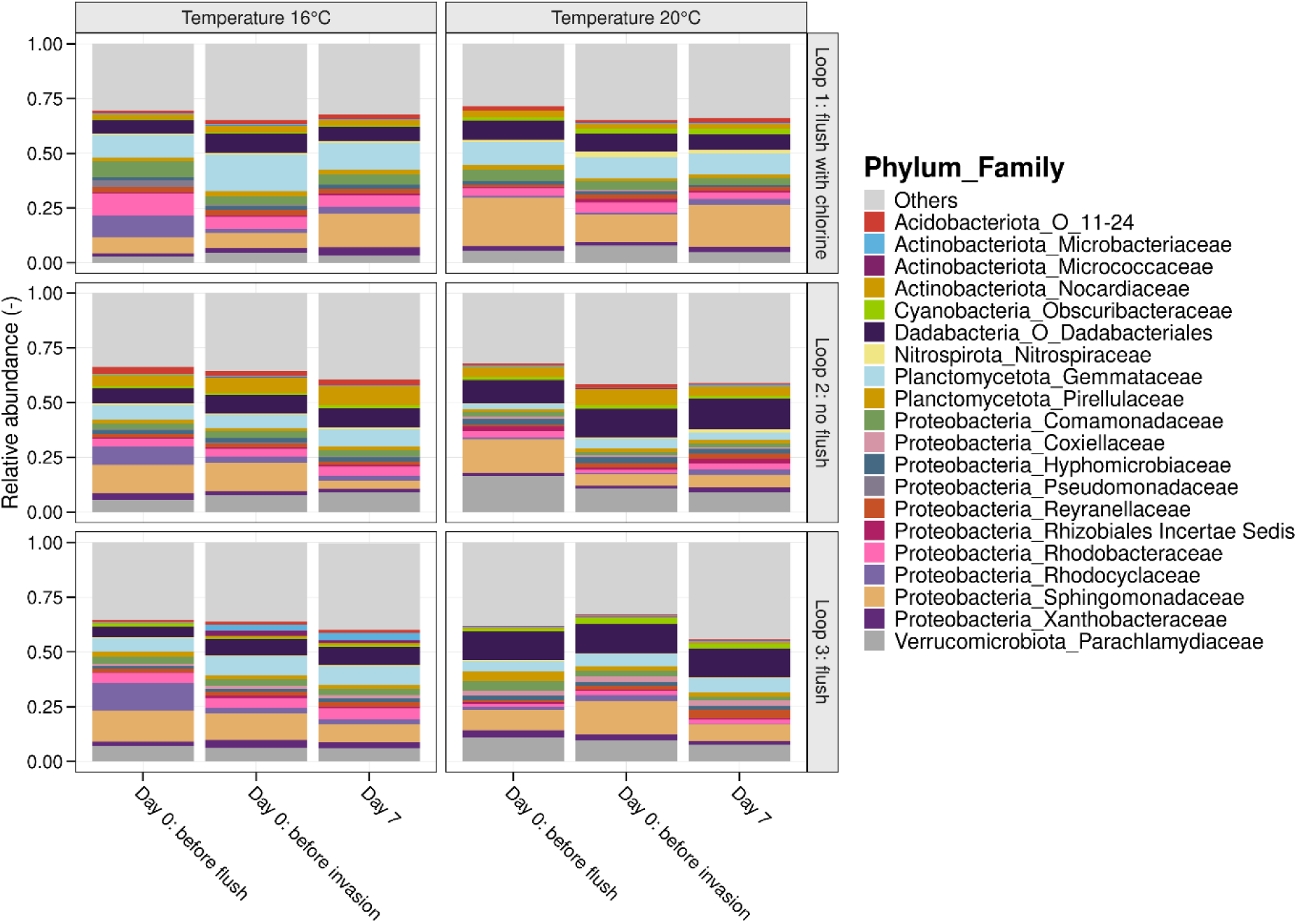
Bacterial community composition in the biofilm. Relative abundances of the 18 most abundant families of the biofilm in each loop before the flush, before the invasion and at the end of the experiment (day 7). Three biological replicates per timepoint were taken during the first experiment (16°C), while the data from the second experiment (20°C) correspond to two biological replicates each. A flush with and without chlorination or no flush had no significant influence on the biofilm community (p > 0.05, PERMANOVA).

Results from the 16S rRNA gene-based amplicon sequencing analysis categorized the samples into three groups: bulk water fed to the pilot, bulk water within the pilot, and biofilm samples (Fig. 4). The biofilm samples exhibited the greatest similarity, even though they were grouped by loop. Corresponding bulk water samples of loop 2 and 3 were grouping together. The bulk water samples fell between the biofilm samples and the water fed to the pilot, indicating the influence of the distribution system and biofilm on the bulk water composition in the pilot.

**Fig. 4:**
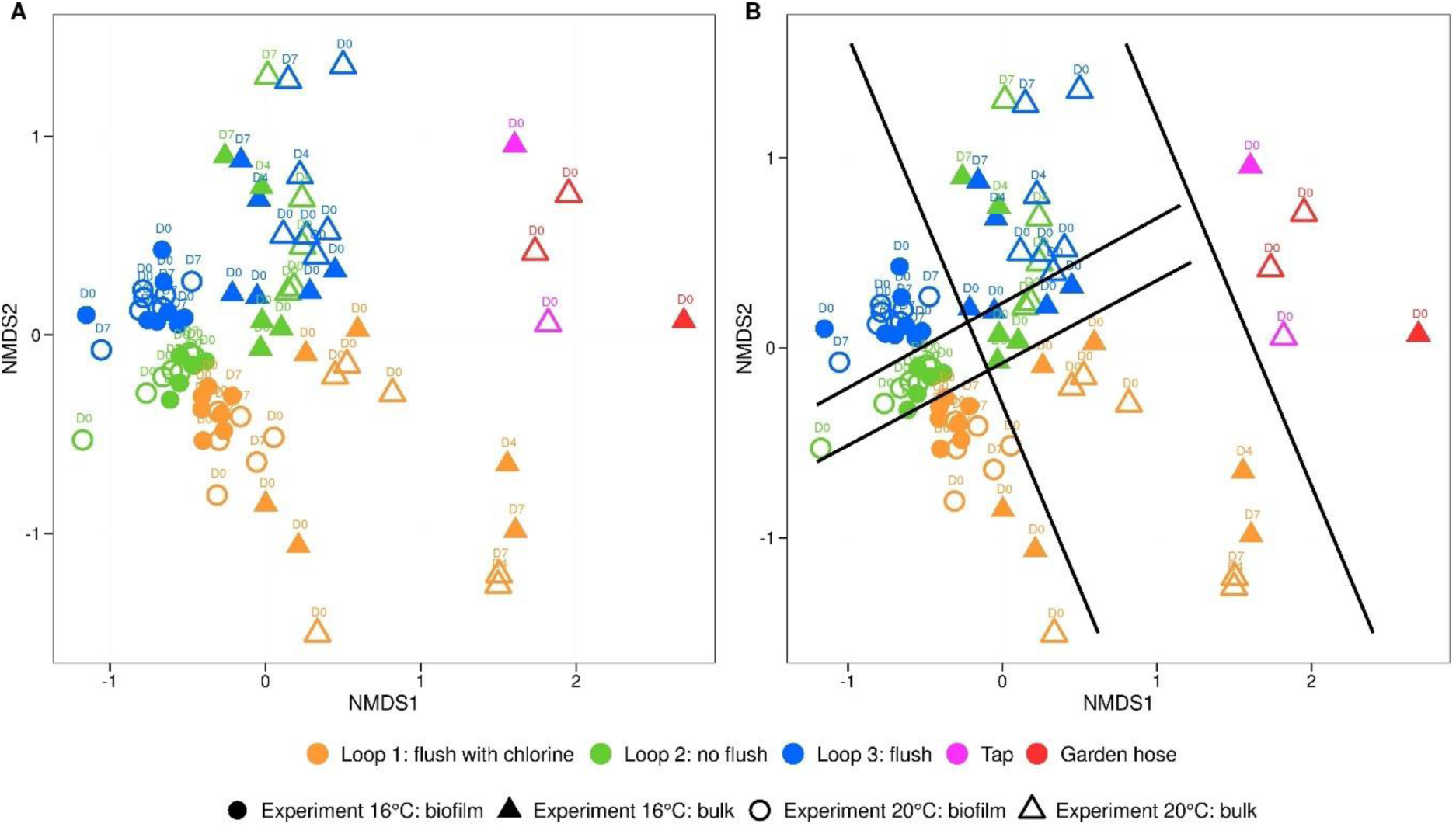
NMDS analysis of bacterial communities in bulk and biofilm samples. NMDS analysis of the relative bacterial community composition (16S rRNA gene) based on Bray-Curtis dissimilarities at amplicon sequence variants levels of bulk (▴) and biofilm (●) samples of loop 1 (flush with chlorine, orange), loop 2 (not flush, green), loop 3 (flush, blue), tap water fed to the pilot (rose) and tap water after the garden hose (red). Timepoints are indicated above each shape. Black lines are dividing the samples in three groups from right to left: the bulk samples of water fed to the pilot, the bulk samples in the pilot and the biofilm samples.

### 3.3 Invasion potential of unwanted microorganisms after flushing procedures

To evaluate the susceptibility of a drinking water network towards invasion after standard operational flushing procedures, an invasion experiment was performed at 16°C and at 20°C. 1 mL of a solution containing 10^6^ cells/mL of *Aeromonas media*, *Pseudomonas putida*, and *Serratia fonticola* was introduced into each loop in order to achieve a final concentration of 100 CFU/100 mL of each microorganism. The survival of the bacterial indicators was followed for seven days using membrane filtration and selective plating techniques. Additionally, first-order decay rate constants (k (h^−1^)) were calculated using Equation 1, with higher k values indicating more rapid decay of the corresponding microorganism. The initial concentrations of *Aeromonas media* and *Pseudomonas putida* ranged from 152 CFU/100 mL to 507 CFU/100 mL and 53 CFU/100 mL to 165 CFU/100 mL, respectively, aligning with the target concentration of 100 CFU/100 mL (Fig. A7). For *Serratia fonticola*, initial concentrations ranged from 242 CFU/100 mL to 400 CFU/100 mL at 16°C, but were between 2070 CFU/100 mL to 3073 CFU/100 mL higher at 20°C. Throughout both spike experiments, and thus at both temperature scenarios, *Aeromonas media* was not detected anymore in each loop 13 hours after the spike, while *Pseudomonas putida* and *Serratia fonticola* persisted for 150 hours (Fig. 5). This was confirmed by the calculated decay rate constants, which were in most cases 10 times higher for *Aeromonas media*, indicating a faster decay compared to the other indicators in drinking water.

**Fig. 5:**
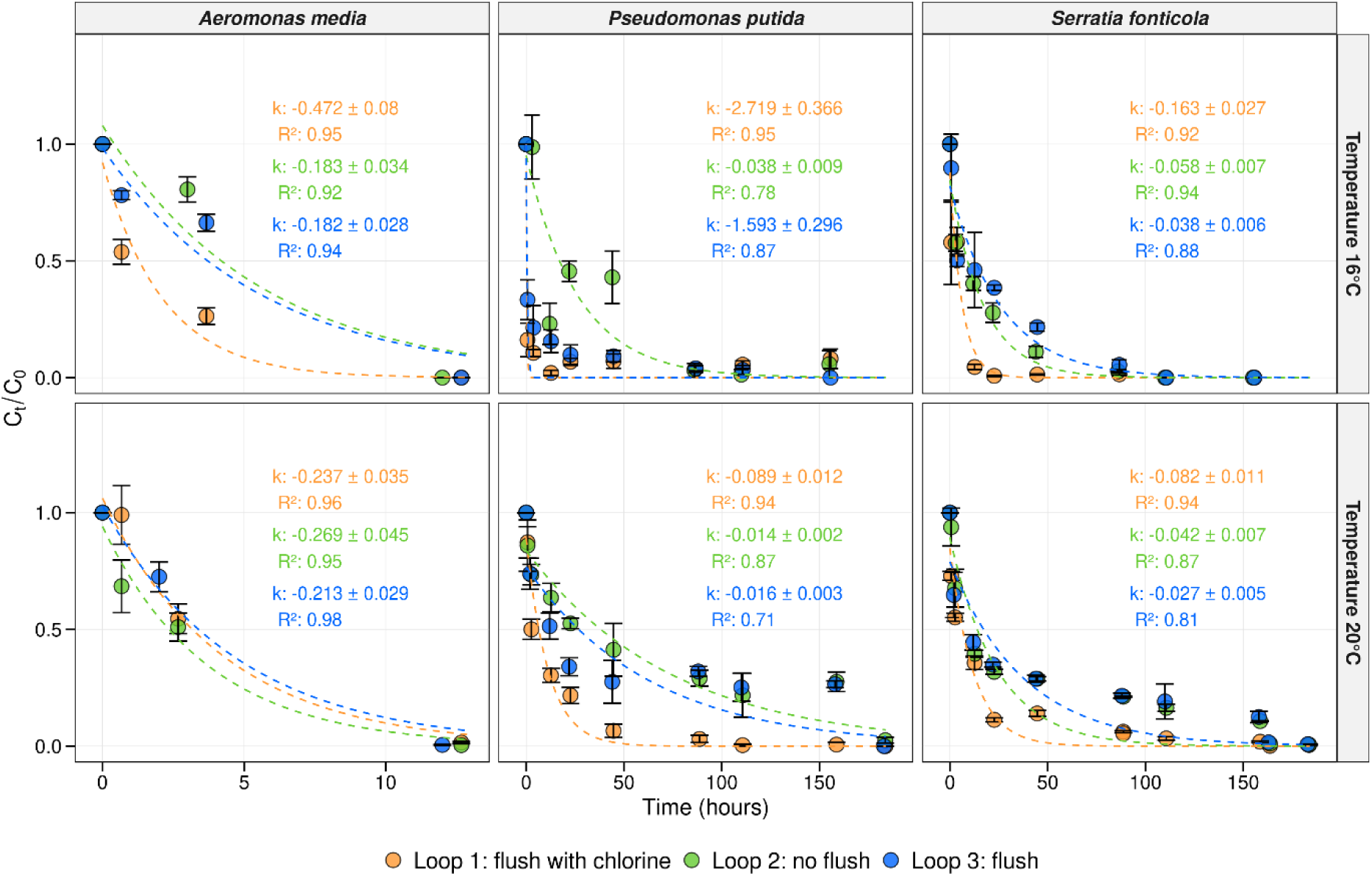
Concentration and decay rates of the bacterial indicators. Average concentration in function of time (hours) for each bacterial indicator at each temperature scenario in loop 1 (orange), loop 2 (green), and loop 3 (blue). The average concentration was calculated as *C*_*t*_/*C*_0_were *C*_*t*_ represents the average concentration at time t, and *C*_0_ the average concentration at the initial timepoint (t = 0). Per timepoint, biological replicates (n = 3) were taken and corresponding error bars are shown in black. First-order decay rate constants (k (h^−1^)) were calculated using Equation 1. The model predictions, along with their respective R squared values, are depicted for each loop with corresponding colors, represented by dotted lines.

At 16°C, the decay rates of *Aeromonas media* were similar in loop 2 and 3 (e.g., 0.183 h^−1^ and 0.182 h^−1^), whereas loop 1, which was flushed and chlorinated, had a slightly higher decay value of 0.472 h^−1^. After 4 hours, a significant difference was observed between the loops (one-way-anova, p < 0.05). At 20°C, decay values of *Aeromonas media* were lower similar across the three loops (0.237 h^−1^, 0.269 h^−1^, 0.213 h^−1^). For loop 1, this value was lower, indicating that higher temperatures led to a slower decay of *Aeromonas media*. After 4 hours, no significant difference was observed between loop 1 and 2 (Tukey test, p = 0.78), while a significant difference was present between loop 3 and the other loops (Tukey test, p < 0.05).

At 16°C, the decay rate constants of *Pseudomonas putida* were similar for loops 1 and 3 (2.719 h^−1^ and 1.593 h^−1^, respectively), and loop 2 had a considerably lower decay rate (0.038 h^−1^). The concentration of *Pseudomonas putida* decreased within 50 hours, but the microorganism remained detectable at low levels (e.g., 2 – 15 CFU/100 mL) in each loop up to 156 hours after the initial spike. At 20°C, both the abundances and decay rate constants showed a more rapid decrease in loop 1 (0.089 h^−1^), which was six times faster than in loops 2 and 3 (0.014 h^−1^ and 0.016 h^−1^). In loop 1, *Pseudomonas putida* was no longer detectable after 155 hours, while it persisted for 184 hours in loops 2 and 3. This was confirmed by statistical tests, which indicated a significant difference between loop 1 and the other loops after 4 hours (Tukey test, p < 0.05), but no significant difference between loops 2 and 3 (Tukey test, p = 1.00). Each microorganism had a slower decay at 20°C compared to 16°C.

Similar trends were observed for *Serratia fonticola* at both temperature scenarios, with a more rapid decay in loop 1 compared to loop 2 and 3, and a more rapid decay at 16°C compared to 20°C (Fig. 5). *Serratia fonticola* was no longer detectable after 111 hours at 16°C and after 184 hours at 20°C. After 4 hours, no significant difference was observed between the loops at both temperature scenarios (Tukey test, p > 0.05), while, after 12 hours the differences between loop 1 and the other loops were significant (Tukey test, p < 0.05). The results showed that both flushing with and without chlorination influenced the decay of the unwanted microorganisms, with the effect being more pronounced at 20°C. This enhanced effect is likely due to competition with the resident drinking water microbiota. In addition, biofilm samples were taken before and after flushing, and again after 7 days, to assess whether any bacterial indicators were able to infiltrate the pretreated biofilm. No unwanted microorganisms were detected in samples taken before and after flushing under both temperature scenarios. For the samples collected after 7 days, no indicators were detected, except for a replicate of the 20°C experiment in loop 2, where *Serratia fonticola* was positively identified, resulting in 5 cells/cm².

## 4. Discussion

### 4.1 Flushing without chlorination does not impact bulk and biofilm cell density and composition

Flushing procedures are widely used by drinking water providers to address quality issues like brown water, or contaminant detection. These processes discharge large volumes of water through hydrants, sometimes with additional disinfectants like free chlorine(9–13). While effective for resolving brown water, flushing could compromise drinking water biostability, increasing the risk of microorganism invasion. This study examined the immediate and long-term impacts of standard flushing procedures (with and without chlorination) in a pilot-scale DWDS with a mature biofilm. Using three identical loops of 100 m PVC pipe, this pilot allowed controlled investigation of bacterial bulk water and biofilm communities under real-life conditions(41). Loop 1 was initially flushed with water containing a free chlorine concentration of 0.50 mg/L. Following this, the loop was flushed twice with water devoid of free chlorine. Loop 3 was subjected to three flushing cycles, while loop 2 acted as a control, with water recirculating continuously for the entire duration of the flushing procedure (7 hours) at a flow rate of 0.08 m/s (shear: 0.03 N/m²) (Table 1). After these flushing procedures, drinking water quality was followed for 2 weeks.

The used flow velocity and shear stress during the flushing procedures (i.e., 0.50 m/s, shear: 0.70 N/m²) were in the same order as the velocities and shear stress used in previous studies in full-scale distribution networks (i.e., 0.11 - 1.00 m/s, shear: 0.03 - 2.64 N/m²)(10, 58) or in pilot systems with PVC/HDPE pipes or pipes imported from the full-scale DWDS (i.e., 0.57 - 1.6 m/s, shear: 0.88 - 7.22 N/m²)(11, 16, 17, 59, 60). On the other hand, lab-scale reactors evaluated higher flow velocities and shear stresses (i.e., up to 10 N/m²) and showed that a shear stress value of 0.2 N/m² can already induce biofilm detachment as biofilm clusters can be affected(18, 61). Although, in practice, the achievable flow velocity primarily depends on the internal diameters, the network’s capacity, and the pressure, meaning our study evaluated relevant flushing conditions(12, 62). Our findings align with previous studies showing that flushing does not fully remove bacteria from pipe walls (Fig. A5), although these studies observed increased turbidity and/or heterotrophic plate counts in the bulk phase, indicating biofilm detachment(10, 15–17, 58, 60). Our study did not detect increased cell densities or TOC concentrations in the flushed water samples (Fig. A2). This may be explained by the cohesivity of the biofilm, which is influenced by factors during growth such as shear stress and substrate material(16, 17). Higher shear stress and turbulent flow as well as smooth pipe materials promote the production of dense, compact biofilms with strong cohesion and a more homogeneous EPS distribution(15–17, 63–65). In our study, flushing was performed on a mature biofilm grown in a controlled environment for 1 year on PVC, a smooth pipe material, under continuous water flow (0.08 m/s). The water residence time was 7 days, with the system being refreshed weekly. During start-up, short bursts of increased flow velocity (0.5 m/s for 5 minutes) were applied, potentially removing the loose surface layers of the biofilm and leaving only the strong basal layer(16). However, employing other methods next to flow cytometry, such as turbidity measurements or more advanced biofilm characterization techniques such as atomic force microscopy, could have provided additional insights. Additionally, we observed no significant impact of flushing on the bacterial biofilm community composition (Fig. 3). Previous studies using young biofilms or mature biofilms imported from the full-scale network reported shifts in bacterial biofilm composition before and after flushing(16, 17). Again, it is important to highlight that the impact of flushing is influenced by a variety of factors(15–17, 63–65). However, our results suggests that mature biofilms with stable density and composition are more resistant to flushing, likely due to their structural stability and cohesive properties.

### 4.2 Flushing with chlorination: no impact on biofilm bacterial cell density and composition but increased cell densities in the bulk over the long term

Sometimes, it is necessary to use an additional disinfectant, such as free chlorine, during the first flushing cycle. The European EPA guidelines recommend disinfection with a local concentration 50 mg/L of free chlorine for 30 minutes, or 20 mg/L of free chlorine for 2 hours, followed by flushing with clean water(9). In practice, lower concentrations are used, with shock dosing applying 10 - 20 mg/L of free chlorine locally, on the spot, for 30 - 60 minutes, in order to achieve 0.25 mg/L residual chlorine throughout the network after 60 minutes of flushing and mixing(12, 18). In our study, free chlorine was dosed in shock to loop 1. After mixing in the loop, a final concentration of 0.50 mg/L of free chlorine was reached and the loop was flushed for 4.3 minutes, resulting in a concentration-time (CT) value of 2.15 mg∗min/L. Following this, the loop was flushed twice with water without free chlorine. Flushing combined with chlorination resulted in an average 1-log reduction in bacterial counts in bulk water (Fig. A2), consistent with the expected bacterial removal by chlorination. For example, Cheswick et al.(66) reported a 1-log reduction using 0.50 mg/L free chlorine for 5 minutes and LeChevallier et al.(67) documented a 99% reduction in the resident drinking water community at a CT value of 3.30 mg∗min/L free chlorine. However, both studies did not incorporate flushing during chlorination, instead using a batch process with stirring. On the other hand, van Bel et al.(11) observed higher removal rates (4 - 6 log reduction) for *Escherichia coli* and *Clostridium perfringens* in bulk water following flushing (1.5 m/s) combined with shock chlorination (10 mg Cl_2_/L, 24 hours).

Before and after the flush with chlorination, no significant impact on biofilm cell concentrations and community composition was observed (Fig. 3, A5). LeChevallier et al.(67) demonstrated that biofilms grown on granular activated carbon were less susceptible than the bulk to free chlorine showing only 25% inactivation at a CT value of 10 mg∗min/L. In addition, a study by Mathieu et al.(18) reported that applying a 60-minute shock chlorination procedure (3.7 mg Cl₂/L) combined with increased hydrodynamic shear stress (1 N/m²) to 2-month-old biofilms led to a 0.7 log decrease of the biofilm cells. These studies used longer contact times and higher free chlorine concentrations on young biofilms, while our study focused on stable biofilms.

Our results showed no direct effect of flushing with chlorination on the cell density and bacterial composition of drinking water biofilms (Fig. 3). However, following the flush, we observed increased bulk water cell densities and a lower phenotypic diversity over the next 12 days, compared to loop 2, which was not flushed, and loop 3, which was flushed without chlorination (Fig. 1). Several studies have reported that disinfection can lead to uncontrolled regrowth during distribution(1, 5, 40, 68, 69). One possible explanation for this is the introduction of dead organic material, which promotes necrotrophic growth of the resident drinking water community(4). Similar microbial families, such as *Comamonadaceae*, and *Pseudomonadaceae*, observed in loop 1 (Fig. 2), were also observed in the necrotrophic grown samples in the study from Chatzigiannidou et al.(4). Additionally, a higher proportion of damaged cells was detected in the biofilm of loop 1 immediately after flushing (Fig. A5B), potentially detaching from the biofilm and contributing to small amounts of nutrients that are responsible for necrotrophic growth. On the other hand, Mathieu et al.(18) showed that shock chlorination (3.7 mg Cl₂/L, 60 minutes) and increased shear stress (1 N/m²) affected the EPS contact points in the biofilm, reducing biofilm cohesiveness. This could have enhanced the release of particles, organic molecules, or dead cells encapsulated within the EPS matrix into the bulk water. Although no significant increase in organic carbon concentrations was detected in loop 1 (Fig. A2), the concentrations may have been below the detection limit of the TOC analyzer (0.03 ppb-C). Various studies have shown that bacteria can grow at very low organic carbon concentrations, even below 10 μg C/L(40, 69, 70).

Notably, prior to the second flush experiment, conducted two months later, elevated cell densities were already detected in loop 1, suggesting that the initial flush with chlorination may have established a new, higher carrying capacity for the resident drinking water community in this loop(1). Additionally, bacteria from the genus *Sphingopyxis* were predominantly growing in loop 1, constituting more than 50% of the community after 7 days. This genus had previously been detected in both bulk water and biofilm at low concentrations and has been identified in several DWDS(71–73). Before the second flush experiment, *Sphingopyxis* was already present at significant levels and exhibited faster growth, reaching over 50% abundance within 4 days. This increase could be attributed to the higher temperature during the experiment (20°C compared to 16°C) or to the fact that *Sphingopyxis* was already more dominant at the start. It is also important to consider that water recirculation in our setup increased the water age, which could enhance the growth of high-abundance groups in the bulk water.

Additionally, we found that the bacterial composition of the flushed bulk water samples (with and without chlorination) was similar to that of water samples collected before the flush (Fig. 4). This is consistent with previous findings that the core microbiome in flushed and tap water samples is largely the same, highlighting the influence of the distribution network on the microbial community composition of bulk water(58). In our research, we showed that the bacterial community in the bulk water samples was more similar to the biofilm than to the water fed into the pilot, emphasizing the importance of distribution pipes and their associated biofilms.

### 4.3 Faster decrease of unwanted microorganisms after flushing with chlorination

To assess the susceptibility of drinking water networks following these flushing procedures, invasion experiments with unwanted microorganisms (i.e., *Aeromonas media*, *Pseudomonas putida* and *Serratia fonticola*) were performed. The invasion experiments were performed with a water temperature of 16°C and 20°C. The start concentration of each indicator in each loop was around 100 CFU/100 mL, 10^5^ times lower than spike concentrations used in previous studies, in order to simulate realistic contamination levels(26, 29, 39, 74–77). The survival of the unwanted microorganisms was followed for 7 days using membrane filtration and selective plating techniques. In general, a decrease of each microorganism was observed over time, with *Aeromonas media* exhibiting a faster decrease compared to *Pseudomonas Putida* and *Serratia fonticola* (Fig. 5). These decreases were confirmed by the first-order decay rate constants calculated as defined by the Chick equation (Equation 1), which were found to be 10 times higher for *Aeromonas media* compared to the other indicators. The decay rates of *Pseudomonas putida* and *Serratia fonticola* were comparable to previously reported decay values of total coliforms and *Escherichia coli* in sewage water and surface water (0.02 - 0.07 h^−1^)(78, 79). Former research showed that *Aeromonas* spp. was not able to compete with the resident drinking water community for nutrients in oligotrophic drinking water, likely due to its preference for biomass components such as amino acids and fatty acids, which are scarce in this environment(69). Additionally, at both temperature scenarios, higher decay rates of *Aeromonas* spp. were observed in loop 2, which was not flushed, while other indicators showed a lower decay in this loop (Fig. 5). This discrepancy may be attributed to the preferential habitat of *Aeromonas* spp. in loose deposits and sediments within the DWDS, that are probably removed in loop 1 and 3 by the flushing procedures(10, 80).

On the other hand, *Pseudomonas putida* and *Serratia fonticola* exhibited a faster decay in loop 1, which had been pretreated with flushing and chlorination. The difference in decay compared to the other loops was more pronounced at 20°C (Fig. 5). The absence of measurable free chlorine indicates that this faster decay was likely due to increased competition, such as for nutrients, because of the higher cell density of the indigenous drinking water community as mentioned before (Fig. 1)(39, 40). Remarkably, *Pseudomonas putida* decreased rapidly within the first 50 hours but remained detectable at low levels (i.e., 2 - 15 CFU/100 mL) in each loop up to 156 hours after the initial spike at both temperature scenarios. This may be due to a more biphasic decay pattern (instead of only exponential), characterized by an initial high first-order rate constant followed by a lower constant, possibly resulting from cells in different growth phases, the presence of different strains, or subpopulations with different resistances to decay(81). Additionally, previous research has reported *Pseudomonas putida* long-term survival and its ability to utilize minimal nutrients in the water, enabling it to persist at low levels despite an overall decrease in numbers(39, 82). Furthermore, *Pseudomonas putida* is known for its resilience and adaptability, allowing it to survive in suboptimal environments(83, 84). Finally, the microorganism may form biofilms or adhere to surfaces within the distribution system, and while not detected by the coupon system, it might be attached elsewhere(85–88).

At higher temperatures (20°C), we observed a lower decay rate for all three indicators, with *Pseudomonas putida* and *Serratia fonticola* also persisting longer in the system (up to 184 hours compared to 156 and 111 hours, respectively) (Fig. 5). These lower decay rates at 20°C align with the idea that warmer conditions can enhance microbial activity and slow down the decay process(89). This finding is consistent with expectations, as higher temperatures often lead to increased numbers of indicator organisms such as coliforms and *Aeromonas* spp.(3, 90) and give a competitive advantage to pathogens like *Escherichia coli* O157(40). Though, the elevated temperatures did not have led to increased growth of the resident drinking water bacteria (Fig. 1). Next to the fact that *Serratia fonticola* persisted over 100 hours, its decay rate was lower compared to other spiked microorganisms, possibly due to the higher initial spike concentration (Fig. A7). After 7 days, *Serratia fonticola* was detected in one replicate of the biofilm from loop 2 at 20°C, the loop that had not been flushed, albeit at a low concentration (i.e., 5 cells/cm²). It is known that drinking water biofilms are heterogenous environments, meaning that certain microcolonies within biofilms may be more susceptible to invasion(91, 92). In a study by Waegenaar et al.(26), *Serratia fonticola* was also detected after 7 days in a mature drinking water biofilm using confocal microscopy and qPCR. In our study, we identified this species through selective plating, which confirms that *Serratia fonticola* was actively present in the biofilm. However, it is important to emphasize that only one replicate showed positive detection, highlighting variability within the biofilm system. Besides, there was no detection of any invader in the biofilms of the flushed loops, indicating that flushing did not lead to more susceptible biofilms for harboring unwanted microorganisms.

In general, we consider the unwanted microorganisms or indicator organisms as r-strategists, characterized by a high growth rate at high nutrient concentrations, whereas the naturally drinking water community consist out of K-strategist characterized by high substrate affinity(1). Our results indicate that the low nutrient concentration of the oligotrophic drinking water environment and the presence of the indigenous drinking water community (possibly through competition for the nutrients) inhibited the spiked bacterial indicators from growing(1, 37, 39, 40). Notably, we introduced the unwanted microorganisms as a one-time contamination event. However, certain unwanted occurrences, such as rainwater intrusions due to improper connections, can lead to periodic or even continuous contamination, potentially enhancing this effect.

## 5 Conclusion

The goal of this study was to evaluate the susceptibility of drinking water networks after standard operational flushing procedures as these procedures can alter biostability. We observed that flushing with and without chlorination did not impact biofilm cell density or community composition. However, flushing with chlorination resulted in increased bulk cell densities and lower phenotypic and genotypic diversity over the long term, indicating that overall biostability was affected. This regrowth can be explained by necrotrophic growth promoted by residual organic material from chlorination or by changes in biofilm structure such as a reduction in EPS contact points. In addition, a decrease of each unwanted microorganism was observed over time in each loop, with *Pseudomonas Putida* and *Serratia fonticola* decaying faster in the loop flushed with chlorination. *Aeromonas media* exhibited a 10 times faster decrease compared to *Pseudomonas Putida* and *Serratia fonticola*. Higher temperatures were found to slow decay, enhancing the persistence of unwanted microorganisms. Our results showed that flushing with chlorination promoted microbial regrowth while accelerating the decay of the unwanted microorganisms, highlighting a trade-off between controlling unwanted microorganisms and potential regrowth.

## Acknowledgments

We would like to thank De Watergroep, Pidpa, and Farys for isolating and delivering the bacterial indicators used in this study. Additionally, we thank Katrien De Maeyer (Pidpa) and her colleagues from the bacteriology lab for performing the MALDI-TOF MS analysis. We also thank Greet Van de Velde (CMET) for conducting the IC measurements.

F.W. is supported by the Research Foundation—Flanders (FWO) (grant number 1S02022N), T.P. is funded by Research Foundation – Flanders (FWO) (grant number 1S26823N) and this study contributes to the FWO-SBO Biostable project (grant number S006221N). The work is part of the Ghent University-Aquaflanders Chair for Sustainable Drinking Water, which is supported by Aquaflanders, the federation of Flemish companies that are responsible for drinking water and sewer management (www.aquaflanders.be).

F.W., T.P. and E.V. carried out the laboratory work, analyzed the data, interpreted the results. F.W. wrote the paper and did additional data analysis. B.D.G. and N.B. interpreted the results and supervised the findings of this work. All authors reviewed and approved the manuscript.

## Data availability statement

The datasets generated and analyzed during the current study are publicly available at https://github.ugent.be/fwaegena/Invasion_Pilot. Flow cytometry files can be sent upon request. Sequencing data were deposited in NCBI SRA (BioProject PRJNA1192987).

